# A novel combination of chemotherapy and immunotherapy controls tumor growth in mice with a human immune system

**DOI:** 10.1101/191031

**Authors:** Aude Burlion, Rodrigo N. Ramos, KC Pukar, Kélhia Sendeyo, Aurélien Corneau, Christine Ménétrier-Caux, Eliane Piaggio, Daniel Olive, Christophe Caux, Gilles Marodon

## Abstract

Mice reconstituted with a human immune system and bearing human tumors represent a promising model for developing novel cancer immunotherapies. Here, we used mass cytometry and multi-parametric flow cytometry to characterize human leukocytes infiltrating a human breast cancer tumor model in immunocompromised NOD.SCID.γc-null mice reconstituted with a human immune system and compared it to samples of breast cancer patients. We observed highly activated human CD4^+^ and CD8^+^ T cells in the tumor, as well as minor subsets of innate immune cells in both settings. We also report that ICOS^+^ CD4^+^ regulatory T cells (Treg) were enriched in the tumor relative to the periphery in humanized mice and patients, providing a target to affect Treg and tumor growth. Indeed, administration of a neutralizing mAb to human ICOS reduced Treg proportions and numbers and improved CD4+ T cell proliferation in humanized mice. Moreover, a combination of the anti-ICOS mAb with cyclophosphamide reduced tumor growth, and that was associated with an improved CD8 to Treg ratio. However, depletion of human CD8^+^ T cells only marginally affected tumor control whereas depletion of murine myeloid cells abrogated the effect of the combination therapy. Altogether, our results indicate that a combination of anti-ICOS mAb and chemotherapy controls tumor growth in humanized mice and highlight the crucial implication of innate immunity in treatment efficacy, opening new perspectives for the treatment of breast cancer.

**One sentence summary:** ICOS expressed on Tregs is a promising target to improve tumor immunity in humans

**Abbreviations:**

ICOS: Inducible Costimulatory
NSG: NOD.SCID.gc-null
Treg: regulatory T cells
CTX: cyclophosphamide
HuMice: humanized mice
CyTOF: cytometry time-of-flight
tSNE: tdistributed stochastic neighbor embedding
pDCs: plasmacytoid dendritic cells
DC: dendritic cells
ICD: immunogenic cell death

## Introduction

Preventing immune suppression in tumors is the “next frontier” in immuno-oncology research. Despite recent success of anti-PD-1 mAbs for metastatic melanomas and other solid tumors, it seems reasonable to propose that combination therapy will be key to the success of cancer immunotherapy, as suggested in patients receiving a combination of anti-PD-1 and anti-CTLA-4 (1). Likely, the most efficient combinations will emerge from the ever growing list of mechanisms that prevent an efficient immune response to the tumor. Recently, a murine study combining a tumor-targeting mAb, a recombinant cytokine, an anti-PD-1 mAb and a T cell vaccine achieved a remarkable efficacy at clearing large established syngeneic tumors (2), illustrating the power of combinations for T-cell and innate cell mediated tumor control. However, studies purely performed in syngeneic models are only useful to gain biological insights but are not relevant to validate mAbs or drugs selectively targeting human cells. Therefore, a mouse model in which the impact of human-specific mAbs on tumor control could be tested would be most invaluable.

It has been shown that human breast tumor morphology and metastatic properties were conserved in xenografted immunodeficient mice (3,4). This was also showed for ovarian cancers (5) and latter confirmed for melanomas (6), suggesting that a large panel of human tumors engrafts efficiently in immunodeficient mice and reproduces features of clinical progression. Xenograft models might be useful to target human tumors but these immunodeficient models are not relevant to test drugs that target the human immune system, the very definition of immunotherapy. Mice carrying a human immune system (HuMice) and human tumors could therefore represent a relevant model for cancer immunotherapy research, at the interface of mice and humans (7). Most of the literature regarding the use of HuMice for immuno-oncology relies on direct transfer of total PBMC into immunodeficient mice of various genetic backgrounds. Therapeutic efficacy of a combination of an anti-PD-1 and an anti-4-1-BB mAbs has been demonstrated in a model in which the tumor and the PBMC were from the same patient (8). However, PBMC-implanted mice invariably and rapidly suffer from Graft vs Host Disease (GVHD), which complicate the interpretation of the results and limits the duration of the experiments. To preclude this problem, NOD.SCID.gamma-c-null (NSG) immunodeficient mice can be reconstituted with human CD34^+^ hematopoietic progenitors (9), most often isolated from cord blood. Because we demonstrated that a pool of human T cells with a diverse repertoire was then generated (10), it became feasible to assess combination therapy targeting human tumors and/or human T cells *in vivo*, generating results in a setting closer to human physiology than syngeneic murine models. However, there is still a lack of information on the composition and the function of human leukocytes infiltrating human tumors in CD34-reconstituted HuMice. Indeed, very few studies have evaluated the nature of human leukocytes in these models (11–15). Activated T cells in the tumor were observed in a peculiar model of tumor implantation at birth in NSG HuMice that did not lead to tumor rejection (11). A more advanced model of HuMice (MISTRG) was used to decipher the contribution of the innate immune system to tumor control, but the presence of T cells in the tumor was not reported (13). Very few T cells were detected in a Patient-Derived Xenograft (PDX) HuMice model of head and neck squamous cell carcinomas (14). Likewise, very few T cells were detected in breast cancer-bearing NSG HuMice with myeloid cells being the subset the most represented in the tumor (12). Finally, a recent study described the presence of Treg in the tumor of NSG HuMice but a precise quantification of T cells and their associated phenotype was not reported (15). Thus, the presence of human T cells in tumors of HuMice is still open to debate. This is a crucial question to be answered if one wants to use the model in a reliable and relevant way. To document this question, we used cytometry Time-of-flight (CyTOF) (or mass cytometry) allowing quantification of the expression of 30 to 40 proteins at the same time at the single cell level (18), a very useful feature when analyzing samples containing a limited number of cells. When compared to the immune landscape of breast cancer patients, several conserved features were observed. Among those, the Inducible COStimulatory molecule (ICOS) emerged in both patients and HuMice as an eligible marker to target Tregs, a crucial T cell subset dampening the anti-tumoral immune response (19). Here, we provide the first proof-of-principle that a combination of chemotherapy and neutralizing anti-ICOS mAb leads to reduced tumor growth in HuMice, opening the possibility to translate this combination into the clinics.

## Results

### T cell infiltrate in the tumors of HuMice

The general design of our study is represented on Fig. 1A. Irradiated NSG newborns were injected in the liver with cord blood CD34^+^ cells. After 16 to 20 weeks, mice were grafted s.c with MDA-MB-231 human breast cancer cells. After 30 to 40 days of tumor growth follow-up, mice were euthanized and the spleen and tumors were analyzed by mass cytometry. The CyTOF panel was designed to include the most common lineage markers for murine and human leukocytes in order to get a global picture of the immune infiltrate, but also incorporated several activation and proliferation markers of human T lymphocytes (Table S1).

**Fig. 1.**
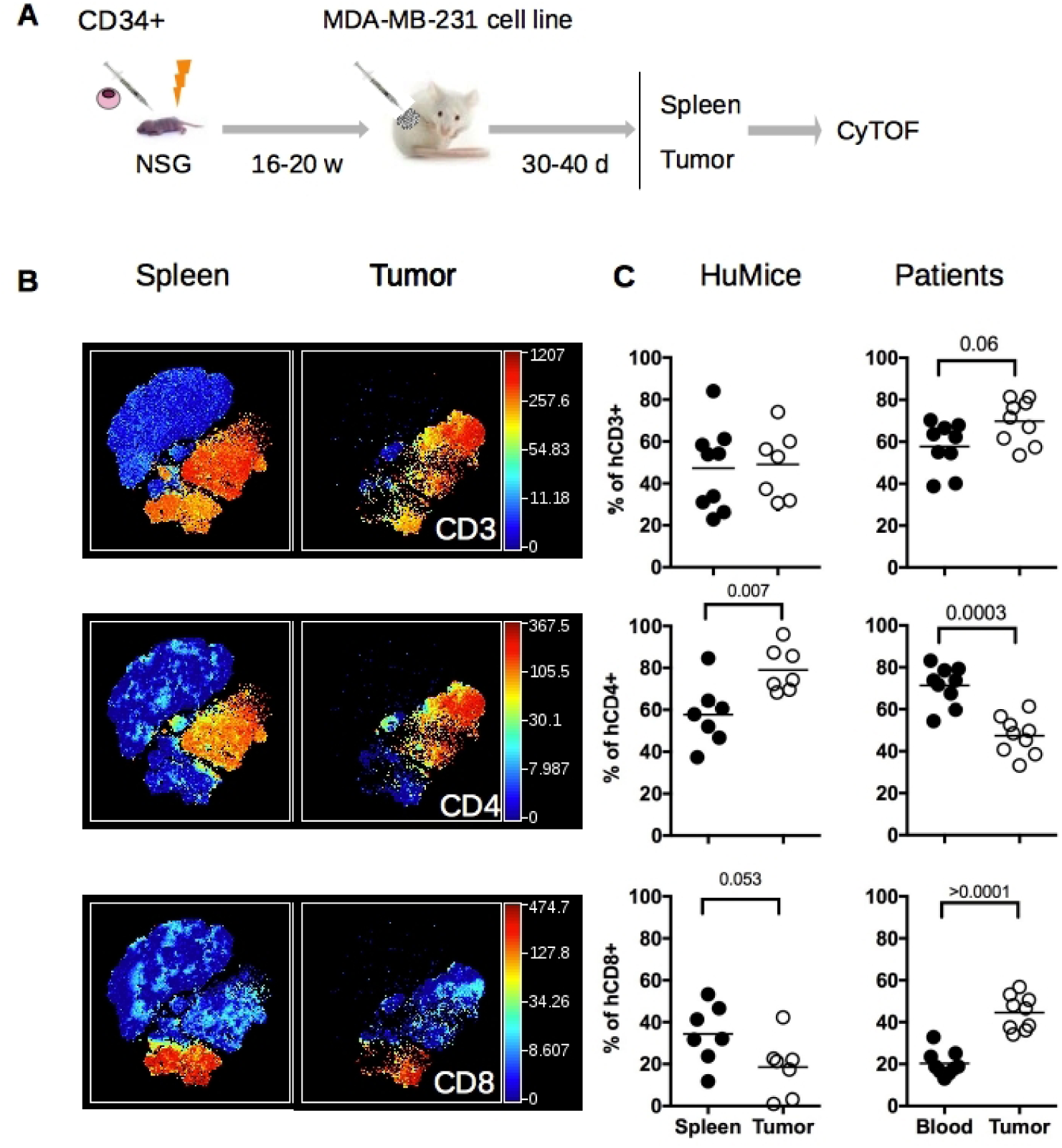
Distribution of human and murine leukocytes in HuMice and in breast cancer patients. **(A)** Experimental scheme of the study, as detailed in the text. **(B)** viSNE representation of gated human CD45^+^ cells in the spleen and the tumor of HuMice from a representative experiment. The viSNE plot was generated according to NKp46, CD38, CD33, CD45RO, PD-1, CD4, CD8, CD20, CD25, Grz-B, and HLA-DR expression using proportional sampling with 54422 events in the spleen and 10274 events in the tumor. The level of expression of each of the indicated markers is represented by a color scale on the right. **(C)** Frequencies of the indicated subsets were determined in HuMice by supervised 2D-gating from CyTOF data and in patients by multi-parametric flow cytometry. Each dot represents a mouse from four independent experiments or a patient in individual experiments. Horizontal line represents the mean value. The p-values indicated in the figures are from non-parametric two-tailed Mann-Whitney t-test.

We first compared the frequencies of human and murine cells of the hematopoietic lineage (CD45^+^) in the spleen and in the tumor in 4 independent experiments, totalizing 15 CD34-reconstituted NSG mice. Within human cells, a classical 2D analysis of the CyTOF data revealed the presence of CD20^+^ B cells and CD3^+^ T cells (CD4^+^ and CD8^+^) in the spleen and the tumor, although B cells tended to disappear from the tumor (Fig. S1A). To get a deeper insight on the nature of immune cells present in the tumor, we performed unsupervised clustering of the data using the tSNE algorithm. This algorithm reduces the multidimensional nature of mass cytometry data to a 2D representation, in which each dot is a cell, clustered according to the level of expression of chosen markers (16). We first focused our analysis on manually gated hCD45^+^ and ran the tSNE algorithm based on lineage and activation markers. Clusters of CD3^+^, CD4^+^ and CD8^+^ cells were readily detected with that method (Fig. 1B). To compare the distribution of human leukocytes in HuMice to human samples, we determined the distribution of human leukocytes in the blood and tumors of 9 breast cancer patients by regular flow cytometry (Fig. 1C). The mean frequencies of CD3^+^ T cells were similar in HuMice and patients in the tumor, representing around 50-60% of total CD45^+^ cells. In contrast, the frequencies of CD4^+^ T cells were higher and the frequencies of CD8+ T cells were lower in the tumor of HuMice compared to patients. Thus, the MDA-MB-231 cell line was indeed infiltrated by human T cells in CD34-reconstituted HuMice, albeit the ratio of CD4 to CD8 T cells was different.

### Innate immune cells in the tumors of HuMice

We next investigated the nature of non-CD19, non-CD3-expressing human cells in the same t-SNE analysis. A distinct cluster of CD33^+^ HLA-DR^+^ cells was observed in the spleen and tumor of HuMice (Fig. 2A). Zooming on CD33^+^ cells, two distinct populations of CD14^+^HLA-DR^lo^ and CD123^+^HLA-DR^hi^ cells further segregated, most likely representing monocytes and plasmacytoid dendritic cells (pDCs) precursors, since pDC precursors and mature pDCs differ by CD33 expression (17) (Fig. 2B). A subset of CD33^+^ cells not expressing CD14 nor CD123, and heterogeneous for HLA-DR were enriched in the tumor, likely including classical DCs (cDC). In addition, two clusters of Granzyme-B-expressing cells (Grz-B^+^) were observed in the spleen and the tumor (Fig. 2C): one expressed CD8 and CD3 (Fig. 1E) and thus represented CTL, while the others expressed CD335 (NKp46) (Fig.2C) but not CD3 (Fig.1D) and thus represented NK cells. In this later cluster, NKp46 expression was lower in the tumor than in the spleen. Thus, in addition to CD4^+^ and CD8^+^ T cells, the immune landscape of tumors in CD34-HuMice also included monocytes, pDC, cDC and NK cells, albeit in small proportions. Similar analysis of human patients was not available.

**Fig. 2.**
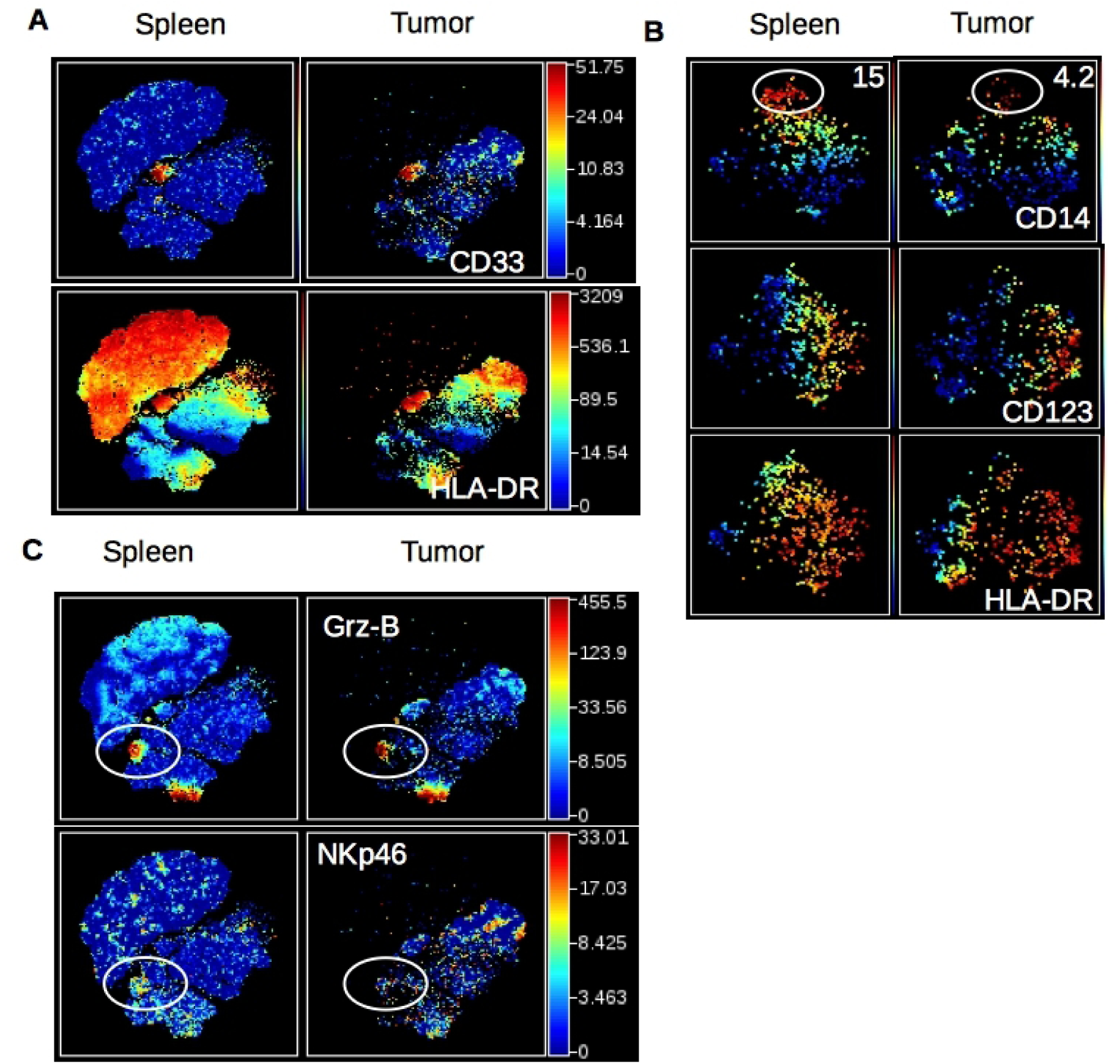
Detection of human myeloid cells and NK cells in HuMice. **(A)** The viSNE plot was generated as in Fig. 1B and represents the expression of CD33 and HLA-DR in hCD45^+^ cells of the spleen and tumor. **(B)** The viSNE plots were generated in gated hCD45^+^CD33^+^ cells according to CD14, CD45RO, CD123, CD4, CD11b and HLA-DR expression using proportional sampling with 535 cells in the spleen and 476 events in the tumors. Indicated on the plots are the frequencies of CD14-expressing cells in the CD33^+^ cluster. **(C)** The visNE plots was generated as in Fig. 1B and represents the expression of Granzyme-B (Grz-B) and NKp46 in the spleen and the tumor. Gates highlight the localization of putative NK cells.

To gain information on the murine infiltrate, the same unsupervised representation of the data was performed on gated mCD45^+^ cells (Fig. S1B). As expected from NSG mice, which are deprived of all lymphoid lineages, most mCD45^+^ cells expressed CD11b in the spleen and the tumor (Fig. S1C). Clusters of Ly-6C^+^ monocytes, Ly6-G^+^ neutrophils and CD11c^+^ DC were readily observed in the spleen and the tumor (Fig. S1C). Overall, myeloid cells of NSG mice were similarly represented in the spleen and the tumor, with the exception of Ly6G^+^ neutrophils which were less abundant in the tumor (Fig. S1D).

### Activated/memory T cells in tumors of HuMice

Having established unambiguously that tumors in HuMice contained human T cells, we next investigated the activation status of those cells. We performed a t-SNE analysis on human CD3^+^ T cells, allowing visualization of two main clusters of CD4^+^ and CD8^+^ T cells (Fig. 3A). When activation/memory and functional markers such as CD45RO, HLA-DR, PD-1, CD25 or Grz-B were considered, we noticed a general increase in the frequencies of cells positive for those markers in the tumor relative to the spleen for both CD4^+^ and CD8^+^ cells, a first indication that T cells were activated in the tumor environment. Up regulation of CD45RO and HLA-DR expression was observed in both CD4^+^ and CD8^+^ subsets whereas higher PD-1 and CD25 expression was noted among CD4^+^ T cells. As expected, up regulation of Grz-B expression was restricted to CD8^+^ T cells (Fig. S2A). To determine the activation status of human T cells more precisely, we performed a boolean analysis calculating the frequencies of cells expressing 0, 1, any combination of 2 or all 3 above mentioned activation markers in CD4^+^ and CD8^+^ T cells. Results depicted in Fig. 3B show that the frequencies of CD4^+^ or CD8^+^ cells co expressing 2 or 3 activation markers were vastly increased in the tumor, while frequencies of T cells expressing none of the activation markers were drastically reduced. Despite this massive activation, the frequencies of proliferating Ki-67^+^ cells co-expressing 3 activation markers were lower in tumors relative to the spleen (Fig. S2B), although this did not reach statistical significance by a 2-way ANOVA. Importantly, a similar enrichment for activated/memory T cells was observed in the tumor relative to the blood in breast cancer patients, assessed by the co expression of 2 or 3 activation/memory markers (Fig. 3C). Thus, tumor infiltrating T cells were of an activated/memory phenotype in both HuMice and breast cancer patients. Of note, PD-1 expression was correlated with higher proliferation of CD4^+^ and CD8^+^ T cells in the spleen of HuMice. In contrast, it was inversely correlated with proliferation of CD4^+^ but surprisingly not of CD8^+^ T cells in the tumor (Fig S2C-E), appearing as an exhaustion marker only for CD4^+^ T cells.

**Fig. 3.**
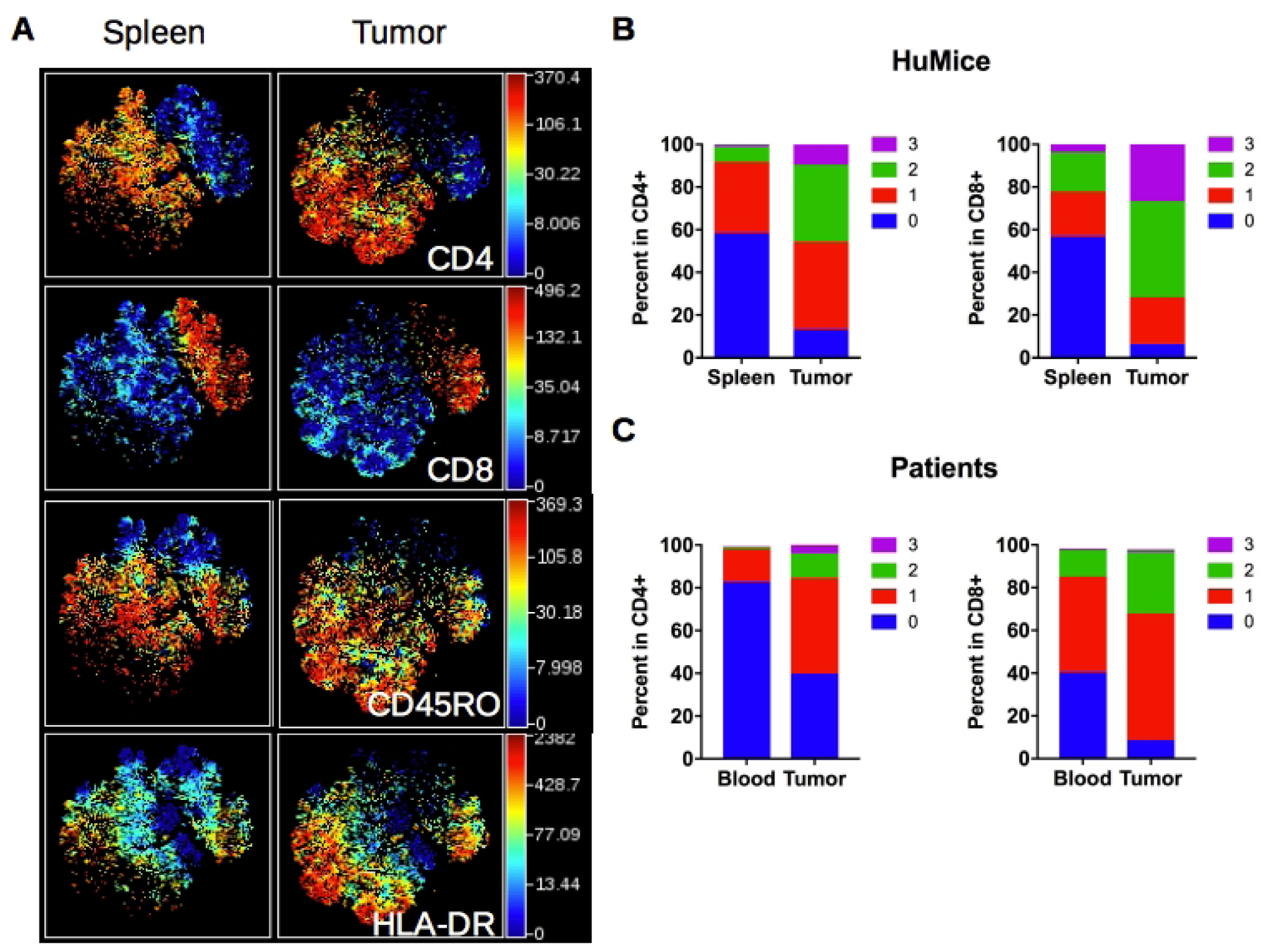
Activation status of Tumor-infiltrating T cells of HuMice and breast cancer patients. **(A)** hCD45^+^CD3^+^ from HuMice were gated in Cytobank and a viSNE plot was generated according to CD4, CD8, CD45RO, HLA-DR, PD-1, CD25, Ki-67 and GrzB expression using equal sampling with 8574 events in the spleen and tumor. **(B)** Mean frequencies of cells expressing 0, 1, any combination of 2 or all 3 activation/memory markers are shown in the indicated tissue. Activation/memory markers used for boolean analysis were CD25, HLA-DR and PD-1 for CD4^+^ T cells and HLA-DR, Grz-B and CD45RO for CD8^+^ T cells. Results are from one experiment out of 3. **(C)** Same analysis for T cells from breast cancer patients. Results are mean frequencies in 9 breast cancer patients in 9 independent experiments in the blood and the tumor determined by flow cytometry. The activation markers used for boolean analysis were HLA-DR, ICOS and PD-1 for CD4^+^ T cells and CD45RA, Grz-B and HLA-DR for CD8^+^ T cells (negativity for CD45RA was considered as an activated/memory phenotype).

### Increased expression of ICOS on regulatory T cells in tumors

Effector Treg are defined by high expression of the transcription factor FOXP3 and the IL-2R-alpha chain CD25, and by the lack of CD45RA expression (18). They are important actors of immunosuppression in the tumors of human patients. It has also been shown that ICOS/ICOS-L interaction can regulate Treg functions in humans (19) and that ICOS expression is linked to Treg-mediated immunosuppression in breast cancer patients (20,21). Furthermore, ICOS expression was associated with poor prognosis in breast cancer patients due to its promoting action on Treg (21,22). A distinct cluster of CD4^+^ cells expressing FOXP3, high levels of ICOS and negative for CD45RA, representing effector Treg was clearly visible in the tumor whereas it was absent in the blood in a patient (Fig. 4A). Across many patients, this cluster was represented at higher frequencies in breast tumors compared to blood (Fig. 4B). It has been shown in humans that CD4^+^FOXP3^+^CD45RA^neg^ cells might include activated effector T cells with lower FOXP3 expression than effector Treg (18). We observed that the frequencies of ICOS^+^ cells were higher in FOXP3^hi^ cells than in FOXP3^lo^ cells in the tumor (Fig. 4C), suggesting that CD4^+^CD45RA^neg^FOXP3^hi^ICOS^+^ cells represented *bona fide* effector Treg. The higher expression of ICOS on Treg of the tumor was also documented by an increase in the MFI of ICOS in FOXP3^+^ vs. FOXP3^neg^ cells (Fig. 4D). Importantly, a similar increase was observed in the tumors of HuMice (Fig. 4D), showing that over expression of ICOS by Treg in human tumors was recapitulated in HuMice. We thus surmise that ICOS could represent a suitable target to affect Treg, with a possible impact on tumor growth, a testable hypothesis in HuMice.

**Fig. 4.**
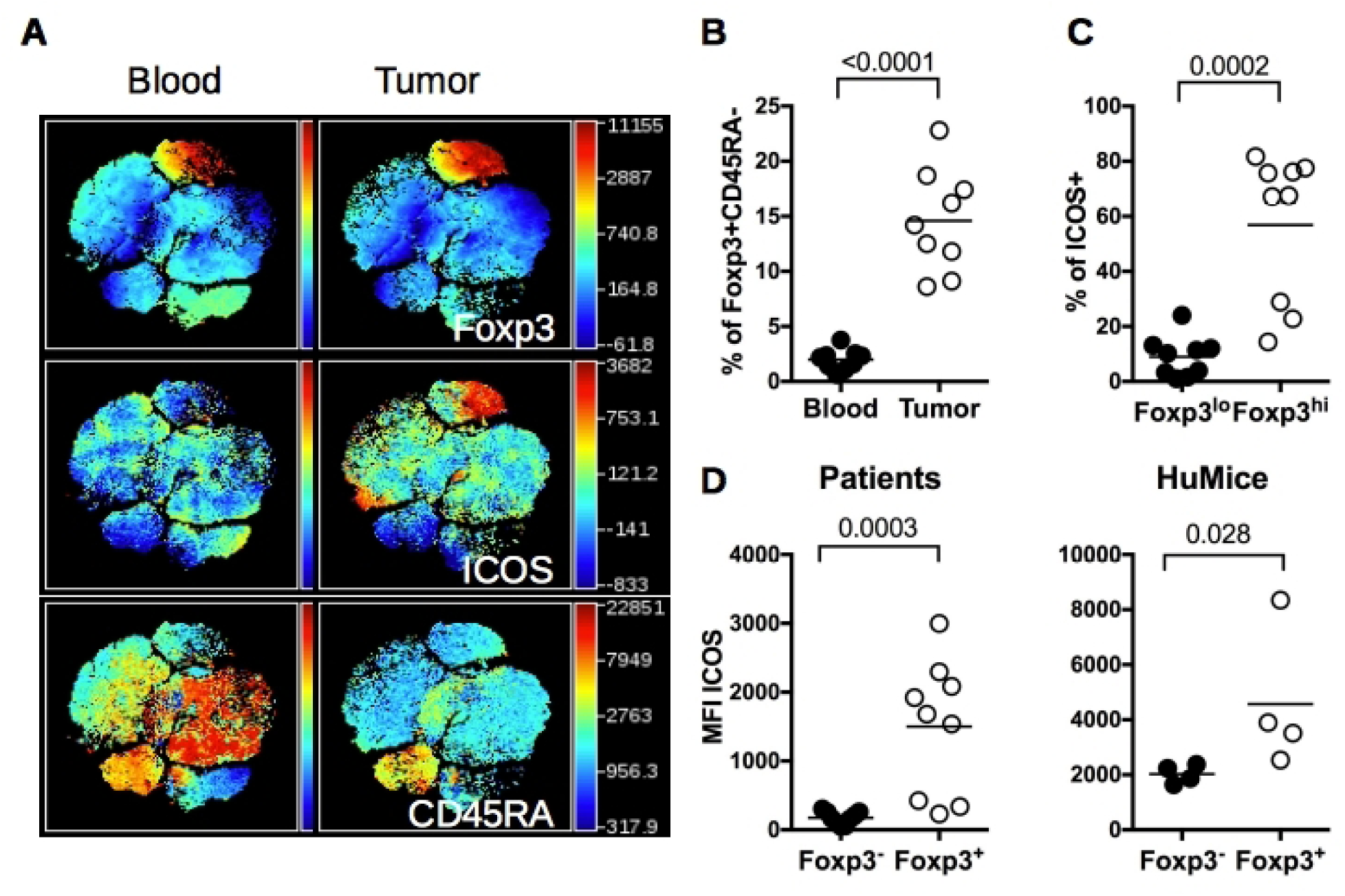
ICOS expression by Treg in HuMice and breast cancer patients. **(A)** viSNE plot of hCD45^+^ from a representative breast cancer patient showing FOXP3, ICOS and CD45RA expression in the blood and the tumor, determined by flow cytometry. **(B)** Frequencies of FOXP3^+^CD45RA^neg^ cells in the indicated tissue from breast cancer patients among CD4^+^CD3^+^ cells. **(C)** Frequencies of ICOS^+^ cells in FOXP3^lo^ or FOXP3^hi^ CD4^+^CD45RA^neg^ in the tumor of breast cancer patients. **(D)** MFI of ICOS in CD3^+^CD4^+^Foxp3^-^ or CD3^+^CD4^+^Foxp3^+^ in the tumor of breast cancer patients or in the tumor of HuMice. Horizontal line represents the mean value. Each dot represents a patient or a mouse. The p values indicated on the graphs are from non-parametric two-tailed Mann-Whitney t-test.

### Impact of anti-ICOS on Treg in HuMice

To test the hypothesis that ICOS/ICOS-L neutralization might affect Treg *in vivo*, we injected once an anti-hICOS mAb, reported as neutralizing *in vitro* (21) in tumor-bearing HuMice and determined the frequencies of Treg in the spleen and the tumor 30 days after (Fig. 5A). Regular flow cytometry was performed to determine frequencies of FOXP3^+^ICOS^+^ and FOXP3^+^CD25^+^ cells following treatment. There was a statistically significant effect of the treatment on the proportions of FOXP3^+^ICOS^+^ cells (p=0.0027, 2-way ANOVA) and of FOXP3^+^CD25^+^ cells (p=0.0039, 2-way ANOVA). By multiple comparisons with corrected p-values, the treatment led to a statistically significant reduction in the frequencies of FOXP3^+^ cells expressing CD25 or ICOS in the spleen of HuMice and similar tendencies were observed in the tumor (Fig. 5A). Moreover, the anti-ICOS mAb treatment was associated with the increased proliferation of total CD4^+^ but not CD8^+^ T cells in the spleen (Fig. 5B). The treatment also affected the absolute numbers of total T cells in the spleen but that did not reach statistical significance. In contrast, Treg counts were significantly reduced by the treatment with numbers dropping almost 20-fold (Fig. 5C). Thus, the anti-ICOS mAb led to a significant reduction of FOXP3^+^ cells in treated animals but also improved proliferation of CD4^+^ T cells, behaving as an antagonist for Treg and an agonist for conventional T cells.

**Fig. 5.**
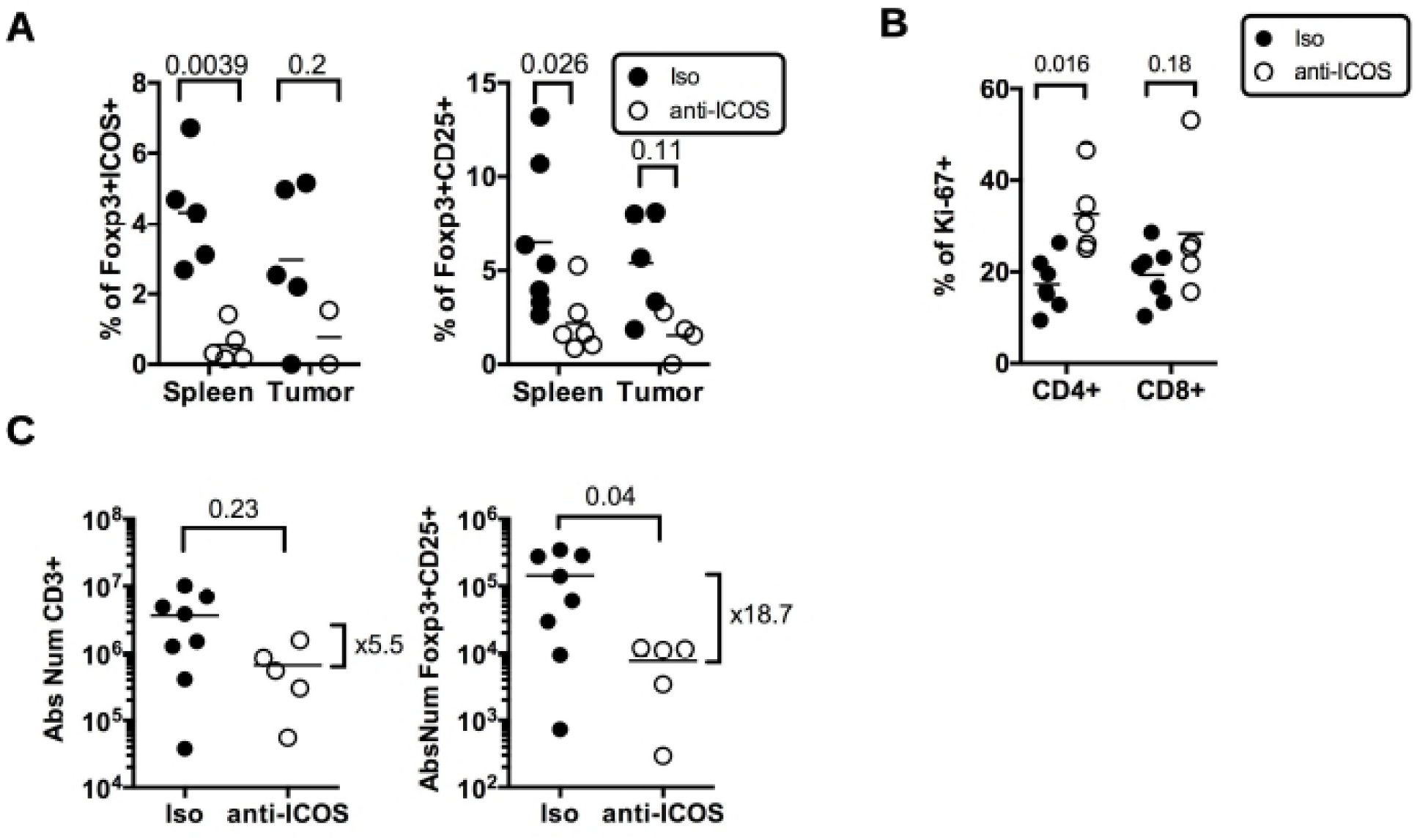
Impact of anti-ICOS mAb on Treg in humanized mice. **(A)** Frequencies of FOXP3^+^ICOS^+^ or FOXP3^+^CD25^+^ cells in CD4^+^CD3^+^ T cells and of **(B)** Ki-67+ cells in CD3^+^ T cells in the spleen and tumor of HuMice injected with isotype control (Iso) or anti-ICOS mAb (50µg/mouse). Results shown are cumulative of at least 3 independent experiments. Only high expressing cells were gated. The p-values reported on the graph are from a 2-way ANOVA with multiple comparisons test corrected by the Sidak method. **(C)** Absolute numbers of total T cells (CD3^+^) or Treg (FOXP3^+^CD25^+^) were based on absolute counts of the spleen of HuMice treated with isotype control (Iso) or anti-ICOS mAb. Each dot represents a mouse. Fold change in mean numbers are indicated on the right. The p-values indicated on the graphs are from an unpaired two-tailed non-parametric Kolgomorov-Smirnov t-test. Data are cumulative of at least 3 independent experiments. Horizontal line represents the mean value.

### Combination of chemotherapy and anti-ICOS to control tumor growth in HuMice

To evaluate the impact of the mAb on tumor growth, we dispatched HuMice with similar T cell reconstitution prior the initiation of the experiments (Fig. S3) into a pre-clinical-like trial with random assignments into groups, blinded evaluation, mixed sex ratio and sufficient number of animals to detect the observed effect with enough statistical power (the experimental scheme is summarized on Fig. 6A). Despite the clear reduction in Treg and CD4^+^ T cells reported above, the anti-ICOS mAb injected alone at day 7 post-tumor implantation had no effect on tumor growth (Fig. 6B). We thus reasoned that combining the anti-ICOS mAb with a known inducer of immunogenic cell death (ICD) might be more efficient, as shown in syngeneic murine models with anti-CTLA-4 or anti-PD-1 mAbs (23). Cyclophosphamide (CTX) is widely used as a chemotherapy for treatment of breast cancer for its cytostatic properties, and has also been described as a potent inducer of ICD and may affect Treg as well (24). Indeed, CTX at a dose of 3 mg per mouse completely abolished tumor growth in non-humanized NSG mice, whereas a dose of 1.5 mg per mouse only moderately affected tumor growth in non-humanized (Fig. S4A) and HuMice alike (Fig. 6B), associated to reduced proportions of Treg in treated animals (Fig. S4B). The combination of the anti-ICOS mAb and CTX injected at day 7 post tumor implantation profoundly reduced tumor growth in HuMice compared to single treatments (Fig. 6B). Thus, a combination of CTX with neutralizing anti-ICOS mAb efficiently controlled tumor growth in CD34-reconstituted HuMice. As expected from the results reported above, the proportion of Treg was lower and the CD8 to Treg ratio was higher in the tumor of the combo group relative to CTX alone (Fig. 6C), showing that the combined effect of CTX and anti-ICOS was associated to a favorable CD8 to Treg ratio.

**Fig. 6.**
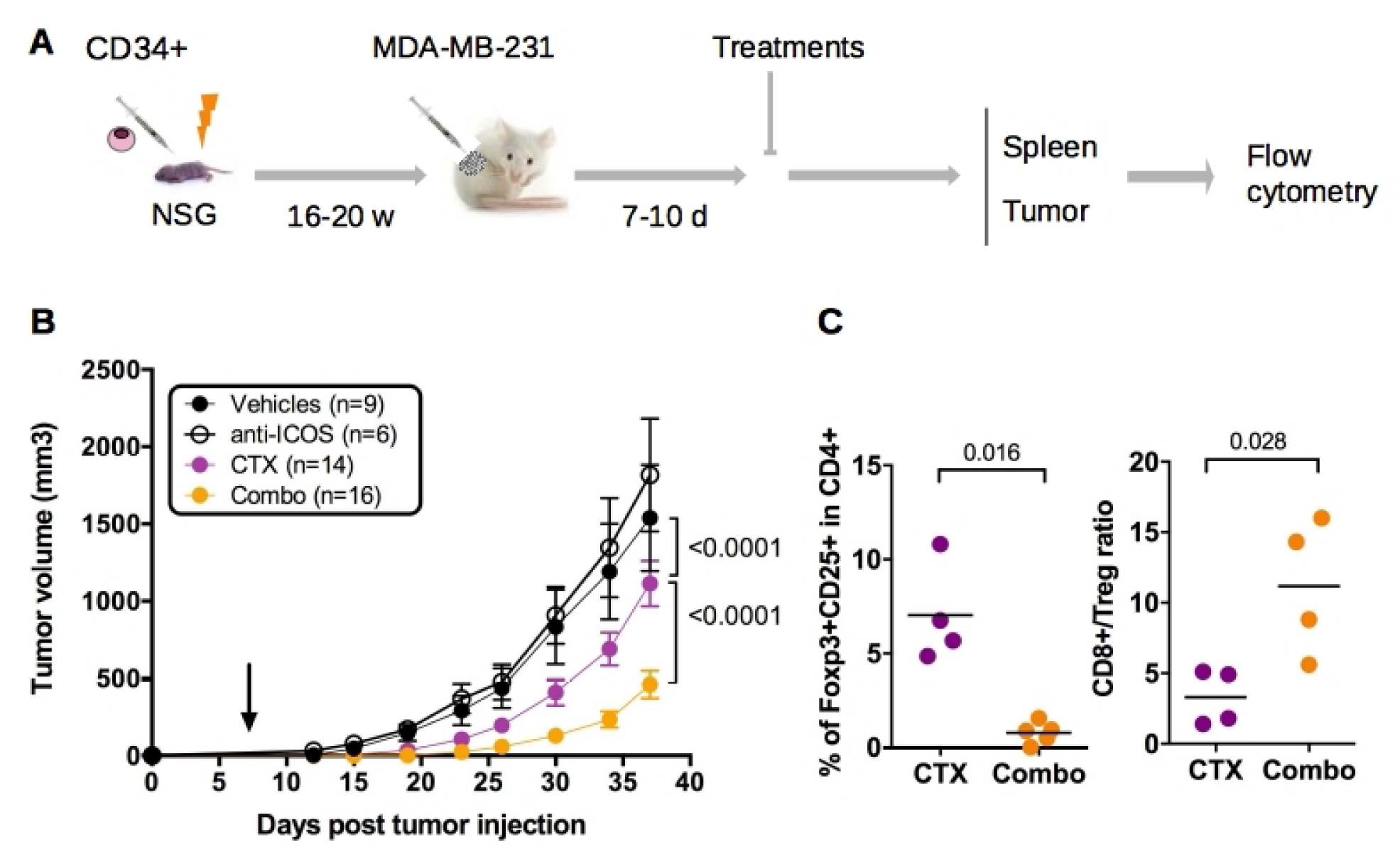
Impact of anti-ICOS mAb on tumor growth in HuMice. **(A)** Experimental design of the study, as detailed in the text. **(B)** Tumor growth was determined in 4 independent experiments in the indicated number of mice treated by PBS and isotype control (Vehicles), anti-ICOS and PBS (anti-ICOS; 50 µg/mouse), cyclophosphamide and isotype control (CTX; 1.5 mg/mouse) or a combination of CTX and anti-ICOS (Combo). Error bars are SEM. The arrow indicates the day the anti-ICOS and the CTX treatment was performed. **(C)** Frequencies of Foxp3^+^CD25^+^ cells in CD4^+^CD3^+^ cells of the tumor in the indicated conditions. The CD8 to Treg ratio was obtained by dividing the frequencies of CD8^+^ T cells by the frequencies of Foxp3^+^CD25^+^ cells in CD3^+^ cells of the tumor in the indicated conditions. Each dot is a mouse and is cumulative from two experiments.

### Role of human T cells and murine myeloid cells in the control of tumor growth by the combination of chemotherapy and anti-ICOS mAb

Frequencies of human CD45^+^CD3^+^ T cells in the tumor were similar in all groups (Fig. S5), indicating that better tumor control in the combo group was not associated to a quantitative increase of human T cells. We thus investigated the role of human CD8^+^ T cells on tumor control in the combo group. For that, we injected HuMice with a chimeric CD8-depleting recombinant Ig (29) after tumor implantation and before the combination of treatments. As already noted in HuMice (30), efficient CD8^+^ T cell depletion was observed in the spleen and tumors of euthanized animals (Fig. S6A) that was quantified by a large increase in the CD4 to CD8 ratio (Fig. S6B). However, tumor growth was marginally affected by the absence of CD8^+^ T cells (Fig. 7A), showing that these were not responsible for better tumor control in the combo group. It was not possible to use a CD4-depleting reagent in HuMice since it would have depleted both Treg and effector CD4 T cells, preventing any conclusions on the sole role of effector CD4 T cells to be drawn. We thus investigated the role of murine myeloid cells on the effect of the combination treatment. We treated a novel set of combo-treated CD34-reconstituted HuMice with the anti-Gr1 mAb that deplete Ly6C^+^ and Ly6G^+^ cells, mostly monocytes and neutrophils. Compared to isotype controls, tumors grew better in the anti-Gr1-treated group (Fig. 7B-C), indicating that murine myeloid cells participated in tumor control evoked by the combination of CTX and anti-ICOS mAb.

**Fig. 7.**
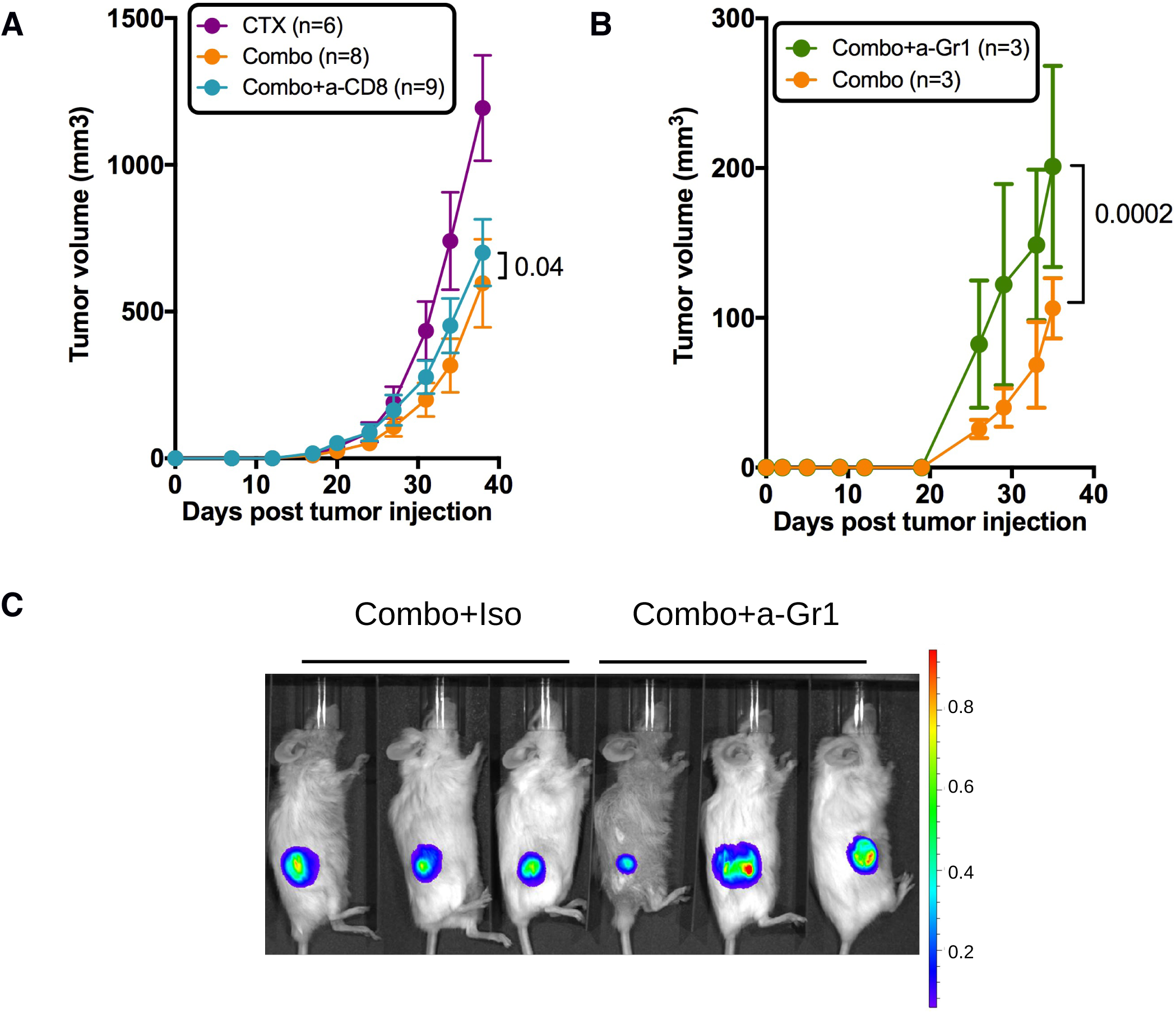
Role of human T cells and murine myeloid cells in the prevention of tumor growth by the combination of chemotherapy and anti-ICOS mAb. **(A)** Tumor growth in the absence of CD8^+^ T cells in the combo group. CD34-reconstituted NSG mice were grafted s.c with the MDA-MB-231 cell line, injected with cyclophosphamide only (CTX; 1.5 mg/mouse) or CTX and the anti-ICOS mAb (50 µg/mouse) with (Combo+a-CD8) or without (Combo) the MT807R1 recombinant Ig (10mg/kg). Results are cumulative of 2 independent experiments (total numbers of mice at the start of the experiment are indicated in brackets). Error bars are SEM. **(B)** Tumor growth was followed in HuMice treated with the combination of CTX and anti-ICOS mAb and an isotype control (Combo+Iso) or an anti-Gr1 mAb (200µg/mouse i.p). The first injection began 2 days before the combo treatment, followed by 5 injections, twice a week for 3 weeks. In this experiment, NSG female mice were humanized at the age of 3 weeks, then grafted as mentioned with the Luc^+^ MDA breast cancer cell line. **(C)** Shown is a representative image 33 days after tumor implantation. The average radiance is represented on a 10^10^ scale on the right. Results are from a single experiment.

## Discussion

In the present study, we provide the first demonstration that CD34-reconstituted HuMice can be used as a platform for the discovery and the validation of novel combinations of chemotherapy and immunotherapy, potentially applicable to patients. This demonstration originates from several similarities between HuMice and patients that we reveal here using multi parametric fluorescent and mass cytometry. A first similarity that we uncovered here was the composition of the tumor immune landscape. Within total human hematopoietic cells, it is remarkable to note that a similar proportion of CD3^+^ T cells infiltrated the tumors in patients and in HuMice. Thus, HuMice models might be useful to decipher the molecules and mechanisms at play that attracts T cells within the tumor environment. We also observed similar infiltration of memory/activated CD4^+^ and CD8^+^ T cells that expressed a combination of activation markers. However, this activation status did not translate into increased proliferation in the tumor of HuMice, suggesting that the tumor microenvironment of this triple negative (TN) PR^neg^ER^neg^HER-2^neg^ cell line might be immunosuppressive *in vivo*. The immunosuppressive status of the tumor microenvironment in breast cancer patients vary according to the nature of the tumor (ER+, HER2+, TN) that may condition T cell infiltrate and T cell activation status (25). Among breast tumor entities, TN are the most infiltrated by T cells (26), CD8^+^ and FOXP3^+^ cells alike, suggesting that TN might be an immunosuppressive-prone microenvironment. Here, we did not compare the proliferative status of T cells from patients with HuMice since none were carrying TN tumors. Further studies should confirm whether the immunosuppressive environment of the TN cell line observed here in HuMice is also observed in patients.

The enrichment for ICOS-expressing Treg is a remarkably conserved feature in HuMice and patients and led us to investigate the therapeutic potential of targeting ICOS for cancer immunotherapy. ICOS, a member of the Ig superfamily, is an essential co-stimulatory molecule for T cell activation and function. Although originally thought of as a co-stimulatory molecule for effective T helper cell response, the biology of ICOS now extends well beyond this narrowed view (27). Relevant to the present study, we and others have shown that the survival, the expansion, and the suppressive functions of human Treg were dependent on ICOS signaling (19,21,28). Thus, we evaluated whether an anti-ICOS mAb could affect human Treg *in vivo*, with possible impact on tumor growth. The reduction in the absolute numbers of total T cells and Treg after anti-ICOS mAb treatment that we report here is in line with an important role for ICOS/ICOS-L on T cells survival. An alternative possibility is that Treg would be physically depleted by the mAb, even though the murine IgG1 isotype used herein is not recognized as a strong inducer of ADCC. This physical depletion is unlikely to occur by complement-dependent cytotoxicity since NSG mice, like their NOD relatives, bears a 2-bp deletion in the C5 gene (29). The murine anti-ICOS mAb might have engaged activating FcRs expressed by myeloid cells, such as FcgRIII, leading to active killing of ICOS-expressing cells, including Treg. Indeed, Treg depletion mediated by myeloid cells has been proposed as a mechanism to explain slower tumor growth following administration of a murine IgG2a isotype to GITR in mice (30). As put forward in the study of Bulliard et al (30), much remains to be done to decipher the mechanisms responsible for mAb-mediated T cell depletion in mice. Whatever the precise mechanisms that remains to be uncover, Treg depletion is now considered a major mechanism for therapeutic efficacy of anti-CTLA-4 or anti-OX40 mAbs in mice (31–33) and possibly of Ipilimumab in humans (34). Additionally, the engagement of ICOS/ICOS-L pathway is required for anti-CTLA-4 efficacy in some models (35). Several mAbs targeting Treg are in development or already approved, including but not limited to anti-CCR4 (36), anti-OX-40 (37), anti-GITR (38) and anti-CD25 mAb (39). In addition, the chemokine receptor CCR8 was recently proposed as an attractive target to affect Treg in breast cancer, although this was not directly demonstrated (40). Our results indicate that ICOS might be a useful addition to this growing list of Treg targets.

In line with published results in murine models (41), we confirm here that blocking ICOS alone was not sufficient to impact tumor growth. However, a combination of chemotherapy and anti-ICOS mAb significantly impacted tumor growth in CD34-HuMice. This result supports the notion that the reduction in Treg is associated to better tumor control if ICD is simultaneously induced by chemotherapy. However, the results of *in vivo* depletion experiments did not support an important role for CD8^+^ T cells on tumor control in the combination group. Thus, we surmise that CD4^+^ T cells might be central for tumor control following the combined treatment. Multiple modes of action for CD4^+^ T cells during anti-tumor immunity are possible and remain to be investigated. Given the results of our myeloid cell depletion experiments, we would like to suggest that CD4^+^ T cells that might have been activated by Treg depletion or directly by the anti-ICOS mAb and that might have attracted murine myeloid cells to the tumor or change their function locally. Supporting this hypothesis, it was recently demonstrated in syngeneic models that combination immunotherapy leads to polarization of monocytes to macrophages (42). This is also remnant of a comprehensive study showing that the proportion of circulating classical monocytes is a good indicator of successful anti-PD-1 treatment for melanoma (43). In addition to human CD3^+^ T cells, our CyTOF analysis showed that the tumors of HuMice also contained rare populations of immune cells known to play important roles during the anti-tumor immune response, such as monocytes or pDCs. The pDC subset that we describe here for the first time in the tumor of HuMice might have played a role on the amplification of ICOS^+^ Treg, as described in breast cancer patients (21). It will be important in the future to monitor the human myeloid compartment more closely in HuMice models optimized for human myeloid development, since those cells can positively (44) and/or negatively affect the local anti-tumor immune response (45). We also revealed the presence of NK cells with low expression of NKp46 in the tumor, suggesting impaired function, in agreement with clinical observations (46).

To our knowledge, our study represents the first demonstration that ICOS represents a target to impact tumor growth in a context of chemotherapy. The presence of ICOS^+^ Treg has been described in breast (20), ovarian (47), and gastric (48) carcinomas, melanomas (49,50) and more recently follicular lymphomas (51), suggesting that ICOS-based cancer immunotherapy might be applicable to a wide range of cancers.

## Methods

### Mice and humanization

NOD.SCID.gc-null mice (stock ≠005557) were originally purchased from the Jackson Laboratory and were bred in our own animal facility under a 14hrs-10hrs light cycle with ad libitum food and water. Mice were given Baytril in their water every other week. Newborns NSG mice were reconstituted by intra hepatic injection of 5. 10^4^ to 10^5^ magnetically-purified CD34^+^ cord blood cells according to the manufacturer’s instructions (Miltenyi, Paris, France) or were purchased from ABCell Bio (Paris, France). Reconstitution was followed over time in the blood by multicolor flow cytometry using the following markers in various combinations of fluorescent dyes: mCD45, hCD45, hCD3, hCD20, hCD4, and hCD8. Validated males and females humanized mice of 16 to 20 weeks old were grafted s.c in the right flank with 1.5.10^6^ MDA-MB-231 breast cancer cells. Mice were euthanized when the tumor reached 3000 mm3 in control groups or 4 to 5 weeks after tumor implantation. All protocols were approved by the French National Ethical Committee on Animal Experimentation (Ce5/2012/025). To assess the effects of the various treatments on tumor growth, a total of 4 experiments are presented. Not every experiment included all conditions. Males and females NSG were randomly dispatched into the various experimental groups to avoid sex-linked effects. Experimental groups were dispatched in different cages to avoid cage-related effects. Tumor growth was monitored in a blinded fashion. The number of mice used in each condition is indicated in the figure legend.

### Cell line

The triple-negative (PR^neg^ER^neg^HER-2^neg^) MDA-MB-231 cell line was grown in DMEM media supplemented with 10% FCS, L-glutamine and antibiotics (Penicillin/Streptomycin) (all from Thermo) in tissue culture flasks. Cells were transduced with a lentiviral vector co-expressing GFP and Luciferase to follow efficient engraftment *in vivo* using luciferase and to allow exclusion of tumor cells from the analysis based on GFP expression. Cells were confirmed of being mycoplasma-free by a standard Hoechst-dye coloration on indicator Vero cells before injection into mice. A genetic profiling was established and confirmed the identity of the cell line (Eurofins Forensic Department, Ebersberg, Germany).

### Reagents preparation and injection

The 314.8 mAb (mouse IgG1 anti-human ICOS) has been described before (21). Isotype controls (mouse IgG1, MOPC-1; rat IgG2b, LTF-2) and anti-Gr1 mAb (rat IgG2b, RB6-8C5) were purchased from BioXcell (West Lebanon, NH, USA). The MT807R1 recombinant Ig consisting of rhesus IgG1k constant regions and CDRs derived from the anti-human CD8 antibody M-T807 grafted into rhesus variable framework regions and was provided by the Nonhuman Primate Reagent Resource (NIH contract HHSN272200900037C and grant RR016001). The antibody was expressed *in vitro* using serum free medium and purified by protein-A affinity chromatography. Endotoxin was <1EU/mg. Cyclophosphamide (CTX, Sigma Aldrich) was prepared extemporaneously according to supplier technical data sheet, i.e to 20 mg/ml of injectable water. All reagents were injected intra peritoneally.

### Phenotypic analysis of leukocytes in the spleens and tumors of humanized mice

Splenocytes and tumors were digested with 1.6 mg/ml of collagenase IV and 10 µg/ml of DNAse I (Sigma Aldrich) for 2 hrs at 37° with regular flushing of the tissue. Cells were passed through a 40µm-cell strainer and resuspended in PBS 3% SVF. To eliminate dead cells and debris, tumor cell suspensions were isolated on a Ficoll gradient. Rings were collected, washed and cell pellets were resuspended in PBS 3%SVF before counting with Trypan blue exclusion. Subsequently, 3 to 6. 10^6^ cells live cells were stained with corresponding antibodies (lanthanide labeled mAbs for CyTOF or fluorochrome-labeled mAbs for FACS analysis). The details of each panel (each one corresponding to one experiment) can be found in Table S1. For CyTOF, 1 to 3µl of each lanthanide-labelled mAbs was added in 1.5 ml Eppendorf tubes in a final volume of 50µl of Max Par Staining Buffer (Fluidigm, San Francisco, USA), according to manufacturer protocol. Intracellular staining was performed with FOXP3 staining kit (eBioscience, Courtaboeuf, France). Cell events were acquired on the CyTOF-2 mass cytometer and CyTOF software version 6.0.626 (Fluidigm) at the Cytometry Pitié-Salpétrière core (CyPS). Dual count calibration, noise reduction, cell length threshold between 10 and 150 pushes, and a lower convolution threshold equal to 10 were applied during acquisition. Data files produced by the CyTOF-2 were normalized with the MatLab Compiler software normalizer using the signal from the 4-Element EQ beads (Fluidigm) as recommended by the software developers. Except for the analysis shown in Fig. 1, GFP exclusion was performed to remove tumor cells from the analysis. To normalize the variability between mice for supervised (i.e 2D plots) and unsupervised (i.e tSNE) analysis, samples from tumors and spleen individually acquired on the CyTOF were aggregated in individual file in each organ and in each experiment.

### Clinical samples

The main clinical characteristics of the patients are summarized in Table 1. Luminal breast tumors were collected from 9 untreated cancer patients undergoing standard surgery at Institut Curie Hospital, in accordance with institutional ethical guidelines and approved by the ethical (CPP ref: 99-15) and medical (ANSM ref: 2015-A00824-45) committees. Flow cytometry data in those patients were collected prior the initiation of the HuMice study, hence no CyTOF data were collected from human patients.

**Table 1.** Clinical characteristics of the primary breast tumors samples. Cod. Pat.= Patient code/ Age= age of the patient at the time of surgery/Meno. status = Menopausal status/ Histo.= Histologic type/ Grade= SBR Grade (histo-prognostic grade based on “Scarff-Bloom-Richardson”)/ pTNM= tumor classification based on Tumor-Nodes-Metastasis score/ ER= % of estrogen receptor positivity/ PR= % of progesterone receptor positivity/ Ki67%= cellular marker of proliferation/ TILs=% of Tumor-infiltrated Lymphocytes.

| **Cod. Pat.** | **Age (years)** | **Meno status** | **Histo.** | **Grad.** | **Size (mm)** | **pTNM** | **ER (%)** | **PR (%)** | **HER2 status** | **Ki67 (%)** | **TIL (%)** |
| --- | --- | --- | --- | --- | --- | --- | --- | --- | --- | --- | --- |
| **808220** | 86 | yes | Lobular | 2 | 21 | pT2N3M0 | 100 | 0 | neg | 20 | 15 |
| **808243** | 55 | yes | Ductal | 2 | 30 | pT2N1M0 | 100 | 15 | neg | 20 | 5 |
| **808254** | 65 | yes | Ductal | 2 | 25 | pT2N2M0 | 95 | 95 | neg | 15 | NA |
| **808364** | 52 | yes | Ductal | 2 | 18 | pT1N2M0 | 90 | 80 | neg | 15 | <10 |
| **808430** | 32 | no | Ductal | 2 | 26 | pT2N2M0 | 100 | 95 | neg | 40 | 20 |
| **809090** | 40 | no | Ductal | 3 | 25 | pT2N2M0 | 90 | 90 | neg | 40 | 30 |
| **809113** | 60 | yes | Ductal | 2 | 28 | pT2N1M0 | 100 | 30 | pos | 17 | 10 |

### Tumor samples from patients and cell isolation

Samples were cut into small fragments, digested with 0.1 mg/ml Liberase TL in the presence of 0.1 mg/ml DNase (Roche, Meylan, France) and incubated for 30 min at 37°C in 5% of CO2 incubator. Subsequently, cells were filtered on a 40-μm cell strainer (BD Biosciences, Le Pont-de-Claix, France), washed and submitted to staining with specific antibodies. Peripheral blood from breast cancer patients was collected in tubes containing EDTA, washed in PBS and submitted to staining with specific fluorescent-labelled antibodies.

### Phenotypic analysis of leukocytes in whole blood and tumors in human breast cancer patients

Tumor cell suspension and whole blood were stained with LIVE/DEAD Fixable Aqua (Life Technologies, Courtaboeuf, France) for 10 min at RT. Then, cells were washed and stained with Aqua BV500 dead cells exclusion dye (Life Technologies); anti-CD3 (clone OKT3, BV650), anti-CD4 (clone OKT4, BV785), anti-PD-1 (EH122.2H7, BV711), anti-CD27 (O323, BV605) from Biolegend (London, UK); anti-CD45 (clone 2D1, APC Cy7), anti-CD8 (clone RPA-T8, BUV395), anti-CD19 (clone HIB 19, Alexa 700) anti-CD56 (NCAM16.2, BUV737), anti-HLA-DR (G46.6, PECy5), anti-PD-L1 (MIHI, PE-CF594) from BD Biosciences; anti-EpCAM (1B7, eFluor660), anti-CD14 (2G5, FITC), anti-CD45RA (HI100, PECy7), and anti-ICOS (ISA3, PERCPe710) from eBioscience for 20 min at 4°C. After incubation, cells were washed and permeabilized for 16 hours according to manufacturer’s instructions for staining with anti-FOXP3 (236A/E7, PE, eBioscience) and anti-Granzyme-B (GB11, BV421, BD Biosciences). Cells were then fixed for subsequent analysis on a Fortessa flow cytometer (BD Biosciences). Data were analyzed with FlowJo Version v10 (FlowJO LCC, Ashland, USA), or Cytobank (www.cytobank.org).

### Mass cytometry data analysis

For CyTOF data analysis, 10 healthy cord blood donors were used in 4 independent experiments, each experiment including mice reconstituted with different donors. A total of 15 mice were analyzed at the steady state by mass cytometry for human and murine cell content in the spleen and the tumor. Due to the paucity of cells in tissues of some HuMice, and to increase resolution of the analysis, tissue samples from 2 to 3 of those mice were pooled before staining and CyTOF analysis. The frequency of hCD45^+^ determined by flow cytometry was used to normalize the representation of each mouse within the pool. In case where sufficient number of cells were collected, tissue samples from individual mice were run into CyTOF. To normalize the analysis from these various conditions, concatenation of individual mice from the same tissue in the same experiment was performed using FlowJo v10. Therefore, this analysis method does not allow comparisons of individual mice but of individual experiments. To document reproducibility of our observations, we provide frequencies of cells in each of the 4 individual experiments, when applicable. Concatenated samples were exported in Cytobank for unsupervised viSNE analysis or were analyzed using FlowJo v10 for supervised analysis. The default settings were applied: 1000 iterations, perplexity of 30 and a theta factor of 0.5.

### Statistical analysis

Statistical analyses were performed using Prism v6.0h for Mac (GraphPad) with statistical models indicated in the figure legends. Outliers detection method is reported in the figure legends when applicable. All tests were performed with an α of 0.05 (probability of detecting a difference in the means by chance below 5%). No a priori sample size estimation to fit statistical power with the observed effect on tumor growth was used. However, a reverse analysis of our data (G-Power; gpower.hhu.de) showed that given the number of mice included in the study, the observed difference in the means at the end of the experiment and the standard deviations in both groups, the β power was >95%, hence validating the rejection of the null hypothesis by statistical modeling of the data. For statistical analysis of tumor growth, the null hypothesis stating that one curve fits all the data in the compared groups was rejected based if the p-value was inferior to 0.05, determined by nonlinear regression modeling of the data using the exponential growth equation.

## Author’s contributions

Conceptualization, AB, CC, and GM; Methodology, AB and GM; Formal Analysis, AB, RR, PKC, KS, AC and GM; Investigation, AB, RR, PKC, KS and AC; Resources, EP, DO and CMC; Writing-Original Draft, AB and GM; Writing-Review and editing, AB, RR, EP, PKC, CMC, CC and GM; Visualization, AC, GM; Supervision, GM; Funding acquisition, CMC, CC, DO and GM.

## Acknowledgements

the authors would like to thank Ms S Just-Landi (Institut Paoli-Calmettes, Marseille, France) for providing the 314.8 mAb, Dr S Brunel for technical help, Dr H. Yssel (CIMI-PARIS, Paris, France) for monitoring mycoplasma infection in cell culture, Dr V. Vieillard (CIMI-PARIS) for advice on NK cells analysis, Dr M. Miyara (CIMI-PARIS) and Dr C. Combadière (CIMI-PARIS) for access to Cytobank, Dr B Salomon (CIMI-PARIS) for critical reading of the manuscript, C. Enond, O. Brégerie and B. Kane (Centre d’Exploration Fonctionnelle, Pitié Salpêtrière, Paris, France) for animal husbandry, and all the mothers for cord blood donations and patients for blood samples.

