## Supplementary material for "A novel combination of chemotherapy and immunotherapy controls tumor growth in mice with a human immune system": Suppllemental

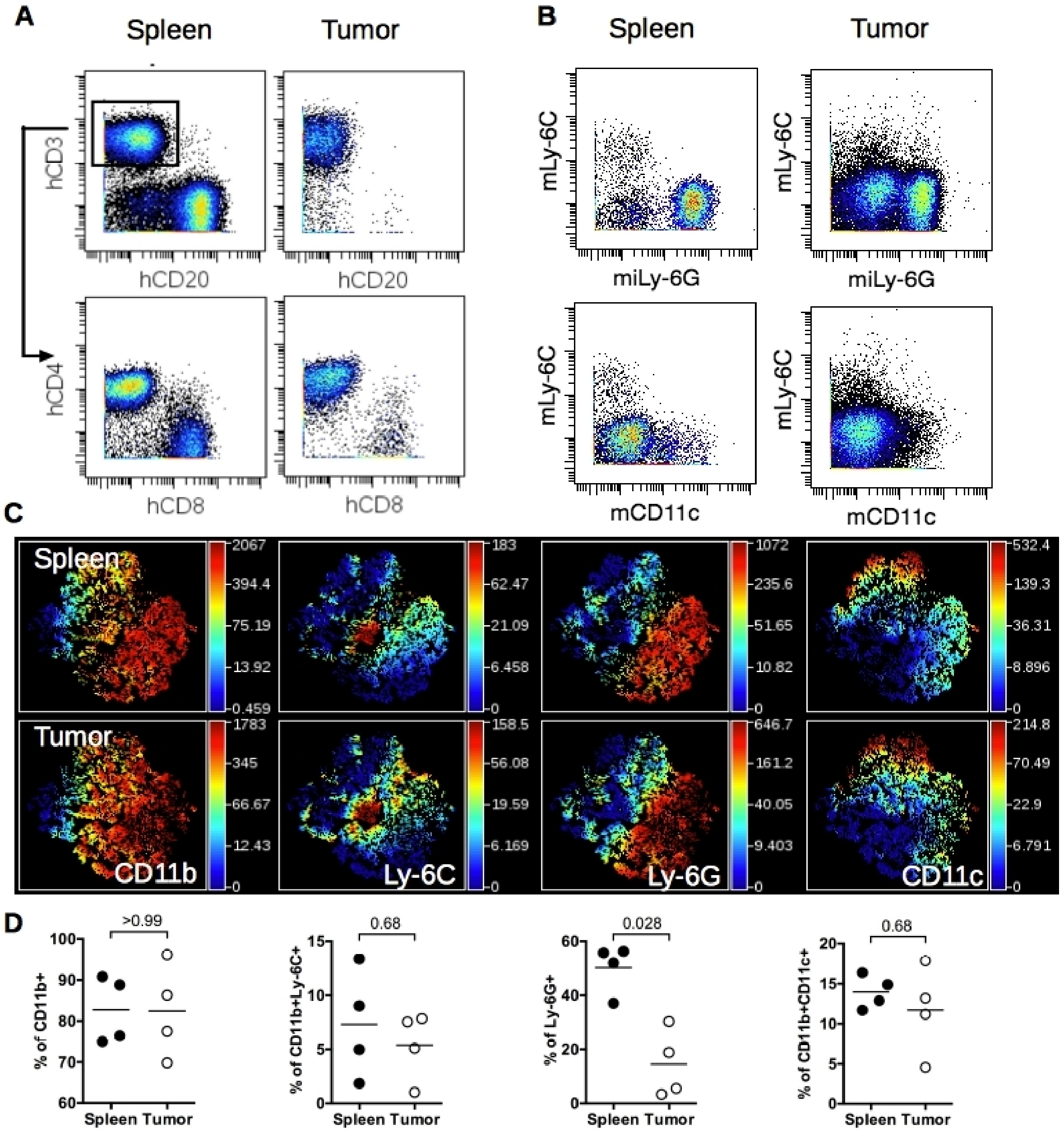


**Fig S1. Distribution of mCD45⁺ cells in spleen and tumors of HuMice. (A-B)** Representative staining in hCD45+ cells (A) and in mCD11b⁺ cells (B) for the indicated markers using CyTOF. **(C)** viSNE representation of gated murine CD45⁺ (mCD45⁺) cells in the spleen (upper panels) and the tumor (lower panels) in a representative experiment. The viSNE plot was generated according to CD11b, CD11c, Ly-6C, Ly-6G expression using equal sampling with 8725 events in spleen and tumor. The plots represent the expression level of the indicated marker depicted by a color code shown on the right. **(D)** Frequencies of mCD45⁺ cells of the indicated phenotype in the spleen and the tumor. Each dot represents an experiment. Horizontal line represents the mean value. The indicated p-values are from non-parametric two-tailed Mann Whitney t-test.

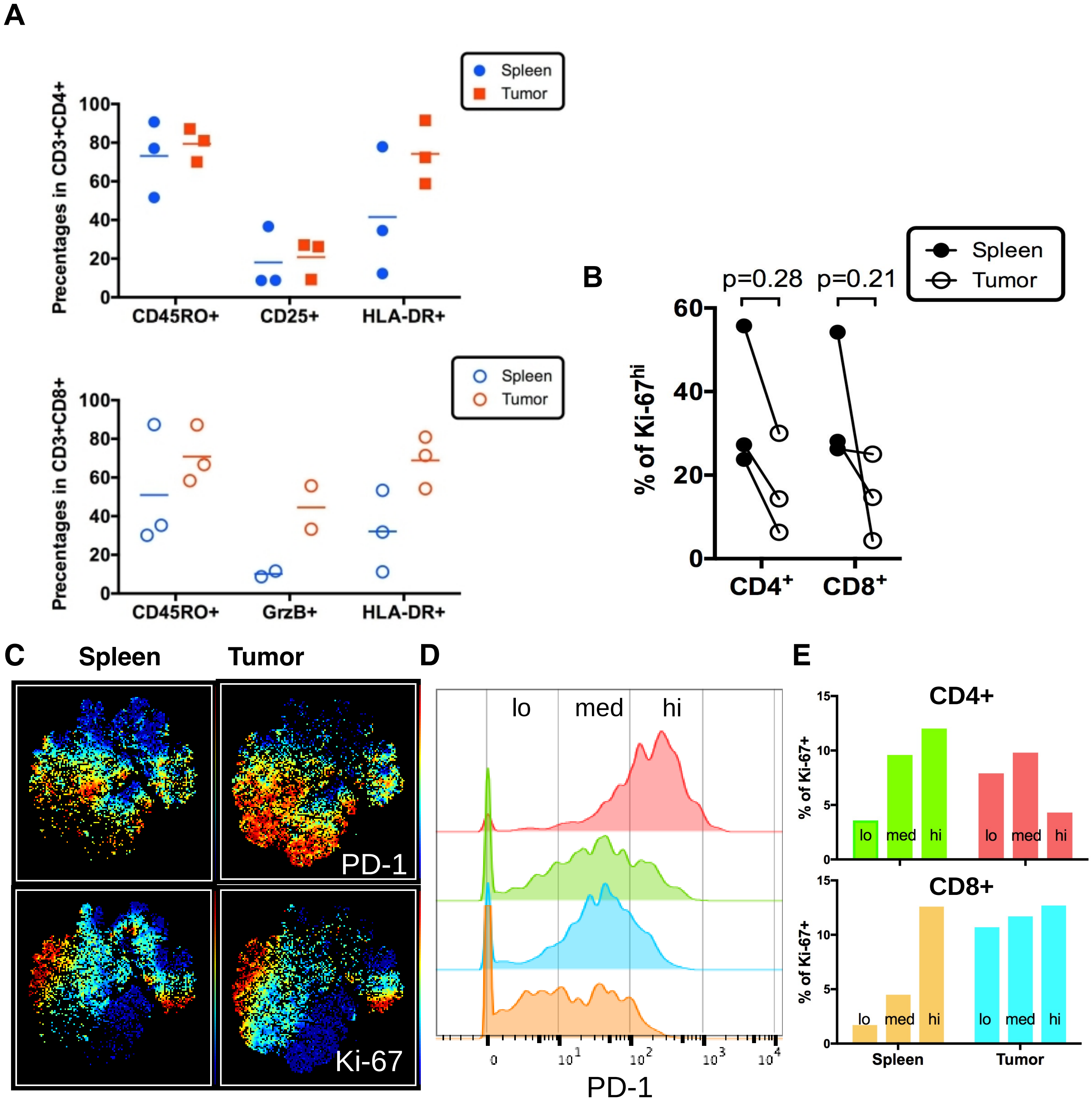


**Fig S2. Proportion and activation status of human T cells in the spleen and tumor of HuMice.** **(A)** Shown are the frequencies of cells expressing the indicated markers in $CD4^{+}$ (upper panel) and $CD8^{+}$ T cells (lower panel) in the spleen and the tumors of HuMice by CyTOF. Each dot represents an experiment. Horizontal line represents the mean value. **(B)** Frequencies of Ki-$67^{+}$ cells in $CD4^{+}$ or $CD8^{+}$ cells co expressing 3 activation markers as in figure 4 in the Spleen or the Tumor of HuMice. Each dot is an experiment. The p-values indicated on the graphs are from a 2-way ANOVA with multiples comparisons using the Sidak correction. **(C)** PD-1 and Ki-67 expression in the spleen and tumor of NSG HuMice without any treatment were determined by CyTOF. ViSNE plots were generated as described in the manuscript. **(D)** Expression levels (low, medium and high) of PD-1 were arbitrarily set in $CD4^{+}$ cells of the spleen (green) or of the tumor (red) and in $CD8^{+}$ cells of the spleen (orange) or the tumor (blue) **(E)** Frequencies of Ki-67+ cells according to PD-1 expression as defined in C.

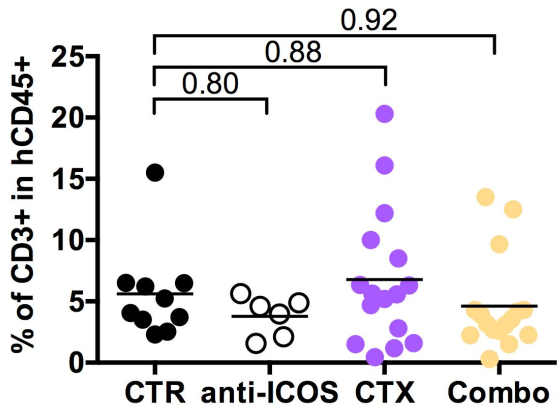


**Fig S3. Frequencies of human T cells in the blood of HuMice prior treatments.** Frequencies of human T cells in the blood prior to the indicated treatments were measured in 12 to 13 weeks-old CD34-reconstituted NSG mice by flow cytometry. None of the treatment groups was statistically different than the control group using one way ANOVA with Sidak's multiple comparisons test. Two outliers were removed from the analysis in the Combo group (ROUT outlier detection, Q=0.5%). Horizontal line represents the mean value.

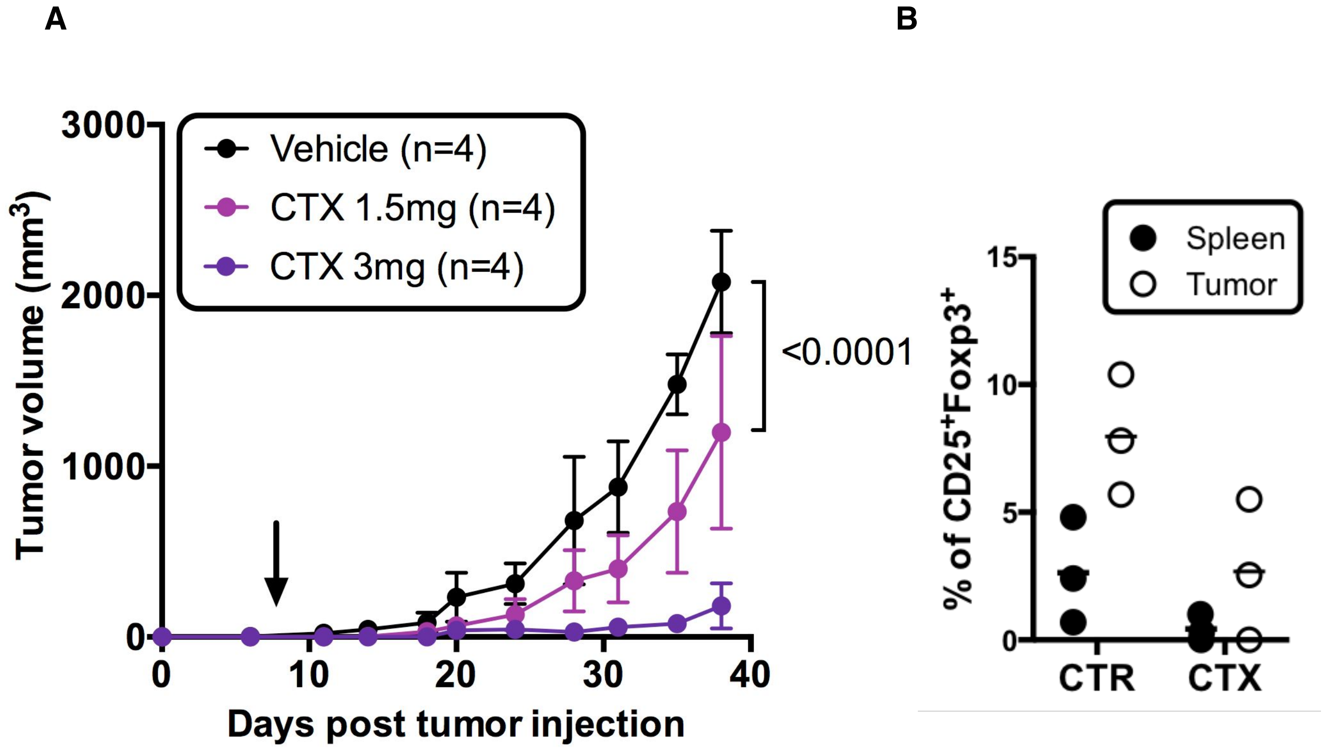


**Fig S4. Effect of cyclophosphamide treatment in NSG and NSG HuMice. (A)** 12 to 16-weeks old females NSG mice grafted s.c with the MDA-MB-231 cell line were treated with the indicated doses of cyclophosphamide (CTX) or PBS (vehicle). Arrow indicates time of CTX injection. The p value indicate the probability that one curve fits all the data between the control (Vehicle) and the 1.5 mg CTX group with a nonlinear regression exponential growth statistical modeling. Error bars are SEM. **(B)** Frequencies of $CD25^{+}Foxp3^{+}$ in $CD4^{+}CD3^{+}$ T cells in the spleen and tumor of NSG HuMice after treatment with vehicle control (CTR) or cyclophosphamide (CTX) were determined by CyTOF. Each dot represents a single experiment including multiple mice.

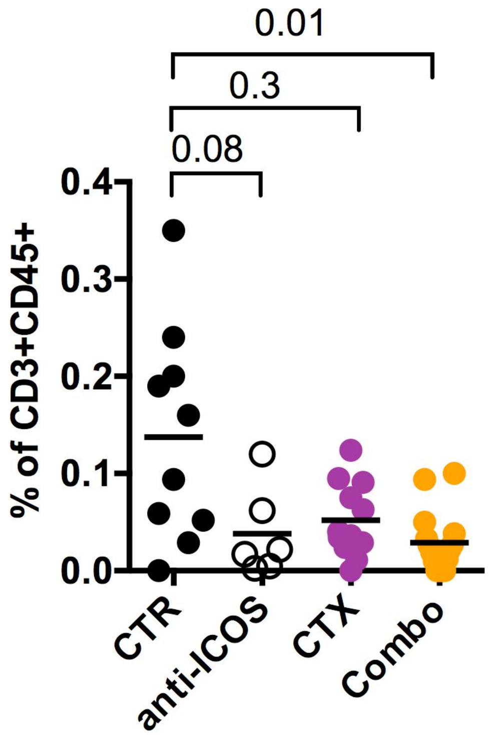


**Fig S5. Proportions of human T cells in the tumor of HuMice.** Frequencies of $CD3^{+}CD45^{+}$ human T cells in the tumor of the indicated groups. Each dot is a mouse. Results are cumulative from 4 independent experiments. One outlier in the CTX and one in the combo groups were detected by the Rout method (Q=0.5%) and were excluded from the analysis. The p-values indicated on the graphs are from a Kruskall-Wallis test with Dunn's corrections.

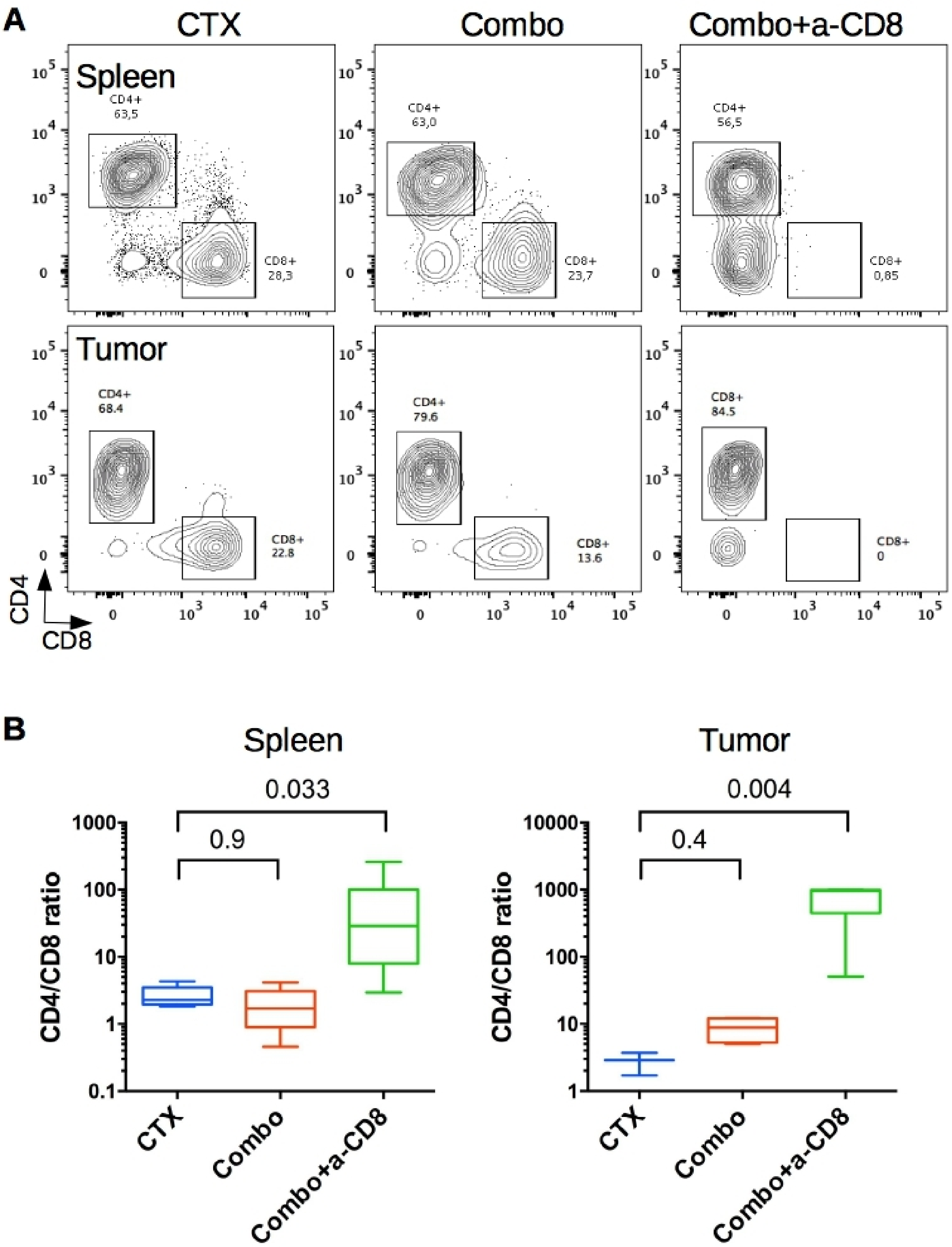


**Fig S6. CD8$^+$ T cell depletion in CD34-reconstituted NSG mice. (A)** Frequencies of CD4$^+$ and CD8$^+$ T cells in the various treatment groups in the spleen and the tumor. Frequencies of each subset are indicated on the graphs. Representative profiles gated in CD45$^+$CD3$^+$ human cells. **(B)** Frequencies determined in (A) were translated into CD4 to CD8 ratio in the indicated groups in the spleen and the tumor. Note the log scale on the left. The p-values are from a Kruskall-Wallis test with Dunn's corrections for multiple comparisons with the CTX group.

**Table S1. Panels used for single cell mass cytometry.**

| Channel | Panel ≠1 | Panel ≠2 | Panel ≠3 |
|---|---|---|---|
| Dy161 | Not used | Not used | hCD49d (P1E6-C5) |
| Dy162 | hFOXP3 (PCH101) | Not used | hCD335 (BAB281) |
| Dy163 | Not used | hCD33 (WM53) | BTLA (MIH26) |
| Er167 | Not used | hCD38 (HIT2) | hCD38 (HIT2) |
| Er168 | Ki-67 (B56) | Ki-67 (B56) | Ki-67 (B56) |
| Er170 | hCD3 (UCHT1) | hCD3 (UCHT1) | hCD3 (UCHT1) |
| Eu151 | hICOS (DX-29) | CD107a (H4A3) | CD107a (H4A3) |
| Gd156 | Not used | CXCR3 (GO25H7) | CXCR3 (GO25H7) |
| Gd158 | Not used | CTLA-4 (BNI3) | hCD33 (WM53) |
| Gd160 | hCD14 (M5E2) | hCD14 (M5E2) | hCD14 (M5E2) |
| Ho165 | hCD45RO (UCHL1) | hCD45RO (UCHL1) | hCD45RO (UCHL1) |
| Lu175 | hPD-1 (EH12.2H7) | Not used | hPD-1 (EH12.2H7) |
| Nd142 | mCD11c (N418) | mCD11c (N418) | mCD11c (N418) |
| Nd143 | hCD123 (6H6) | hCD123 (6H6) | hCD123 (6H6) |
| Nd145 | hCD4 (RPA-T4) | hCD4 (RPA-T4) | hCD4 (RPA-T4) |
| Nd146 | hCD8 (RPA-T8) | hCD8 (RPA-T8) | hCD8 (RPA-T8) |
| Nd148 | Not used | mCD11b (M1/70) | mCD11b (M1/70) |
| Nd150 | mLy-6C (HK1.4) | mLy-6C (HK1.4) | mLy-6C (HK1.4) |
| Pr141 | mLy-6G (1A8) | mLy-6G (1A8) | mLy-6G (1A8) |
| Sm147 | mCD45 (30F11) | hCD20 (2H7) | hCD20 (2H7) |
| Sm149 | hCD25 (2A3) | hCD25 (2A3) | hCD25 (2A3) |
| Sm154 | hCD45 (HI30) | hCD45 (HI30) | hCD45 (HI30) |
| Tb159 | Not used | hFOXP3 (259D) | hFOXP3 (259D) |
| Tm169 | GFP (5F12.4) | GFP (5F12.4) | GFP (5F12.4) |
| Y89 | Not used | mCD45 (30F11) | mCD45 (30F11) |
| Yb171 | hCD20 (2H7) | hCD68 (Y1/82A) | hCD68 (Y1/82A) |
| Yb172 | mCD11b (M1/70) | Not used | Casp3 (5A1E) |
| Yb173 | Not used | Grz-B (GB11) | Grz-B (GB11) |
| Yb174 | HLA-DR (L243) | HLA-DR (L243) | HLA-DR (L243) |
| Yb176 | hCD56 (HCD56) | Not used | Not used |

Indicated in the table are the molecules (h=human, m=murine) targeted by lanthanide-labeled mAbs (clones are indicated in brackets). mAbs highlighted in red did not give any signal over background.
